# Tetraspanin immunocapture phenotypes extracellular vesicles according to biofluid source but may limit identification of multiplexed cancer biomarkers

**DOI:** 10.1101/2021.03.02.433595

**Authors:** Rachel R. Mizenko, Terza Brostoff, Tatu Rojalin, Hanna J. Koster, Hila S. Swindell, Gary S. Leiserowitz, Aijun Wang, Randy P. Carney

## Abstract

Tetraspanin expression of extracellular vesicles (EVs) is often used as a surrogate for their general detection and classification from background contaminants. This common practice typically assumes a consistent expression of tetraspanins across EV sources, thus obscuring subpopulations of variable or limited tetraspanin expression. While some recent studies indicate differential expression of tetraspanins across bulk isolated EVs, here we present analysis of single EVs isolated using various field-standard methods from a variety of *in vitro* and *in vivo* sources to identify distinct patterns in colocalization of tetraspanin expression. We report an optimized method for the use of antibody-capture single particle interferometric reflectance imaging sensing (SP-IRIS) and fluorescence detection to identify subpopulations according to tetraspanin expression and compare our findings with nanoscale flow cytometry. Using SP-IRIS and immunofluorescence, we report that tetraspanin profile is consistent from a given EV source regardless of isolation method, but that tetraspanin profiles are distinct across various sources. Tetraspanin profiles as measured by flow cytometry do not share similar trends, suggesting that limitations in subpopulation detection significantly impact apparent protein expression. We further analyzed tetraspanin expression of single EVs captured non-specifically, revealing that tetraspanin capture can bias the apparent multiplexed tetraspanin profile. Finally, we demonstrate that this bias can have significant impact on diagnostic sensitivity for tumor-associated EV surface markers. Our findings may reveal key insights into the complexities of the EV biogenesis and signaling pathways and better inform EV capture and detection platforms for diagnostic or other downstream use.

## Introduction

The promise of extracellular vesicles (EVs) for unbiased diagnostic and therapeutic application requires identification of ubiquitous markers that can distinguish EVs from the contaminating background of free protein aggregates, microparticles, lipoprotein, and numerous other nanoparticulate assemblies found in human biofluids. The group of tetraspanins CD9, CD63, and CD81 are the most common EV-associated markers reported in the literature and have been used for “total” EV capture in many studies, including ELISA^1,2^, flow cytometry^3,4^, and lab-on-a-chip assays^5,6^. Each of these tetraspanins has been demonstrated to play an active role in EV biogenesis or cargo sorting, suggesting their essential role in the EV secretory pathway^7,8,9^. While initial work suggested the universal enrichment of these membrane proteins in EVs isolated across cell types^10,11^, recent studies show they are in fact heterogeneously expressed across EVs^12,13,12,14^. Such variance in expression suggests the existence of distinct EV subpopulations with significant functional differences. Using this panel of tetraspanins as general “EV-specific” biomarkers may erroneously bias results. To our knowledge, no group has studied the frequency and colocalization patterns of tetraspanins on single EVs across sources and isolation methods, a crucial step in understanding their role as generic EV biomarkers and their colocalization with clinically relevant markers.

There is a lack of techniques that can assess multiplexed expression of membrane proteins at single vesicle resolution across the size range of EVs, with the majority of EVs being between ~30 and 100 nm in diameter^15^. Negative staining transmission electron microscopy (TEM) and cryo-EM provide indirect imaging of protein expression via antibody-bound metal nanoparticle probes, thus are close to ground truth^16,11,17,18^. Yet, it remains technically demanding to produce high quality EM data, and artifacts in sample preparation and imaging projection can affect interpretation. Furthermore, multiplexing is limited to only a few markers at a time, since distinct sizes of metallic probes are needed^16^. Bead-based flow cytometry permits multiplexed protein expression analysis of EVs, but relies on bulk analysis of at least thousands of EVs, and is biased according to choice of bead-bound biorecognition element^4,19^. Commercial flow cytometers can assay protein expression at single EV resolution^20,21^ but only above a critical nominal size of ~80 nm, since detection is limited by particle scattering, thus is dependent on particle diameter, particle refractive index, the angle of collection, and the intensity and wavelength of the laser^20^. Some studies have reduced this size limit by detecting particles using fluorescence instead of scattering, yet are still limited in sensitivity to EVs that highly express a single protein^22^.

Single particle interferometric reflectance imaging sensing (SP-IRIS) combined with antibody-based microchip capture and fluorescence detection represents an emerging technique that can identify single vesicles with a lower size limit of detection compared to scattering based techniques^23^. SP-IRIS has a theoretical detection limit down to single molecule size, but is more practically ~50 nm for routine measurement.^24^ In addition to breaching the diffraction limit, this technique can identify multiplexed expression of membrane proteins on single EVs when paired with fluorescence microscopy and an antibody-fluorophore sandwich assay. A commercial SP-IRIS platform, the ExoView R100 (NanoView Biosciences), utilizes a microarray chip for antibody capture against the tetraspanins CD9, CD63, and CD81 (custom arrays are also available).

In this study, we apply the ExoView platform to examine tetraspanin profiles of single EVs from a variety of sources, including cancer cells, stem cells, and human serum, isolated by either ultracentrifugation (UC) or size exclusion chromatography (SEC). We report that tetraspanins are not homogeneously multiplexed across all EVs, but that consistent multiplexed subpopulations are present in a given EV isolate. We explore the impact of heterogeneous tetraspanin colocalization as a form of bias that may impact sensitivity to specific disease markers, especially for platforms where CD9, CD63, or CD81 capture is the common first step to enrich EVs for downstream biomarking. To examine this, we developed a custom assay on the ExoView platform utilizing non-specific biotin-based EV capture to examine the bias of direct tetraspanin capture in selecting specific subpopulations. This work describes the first comparison of single-EV protein expression by immunofluorescence across a variety of EV sources and isolation techniques and could be applied to identify a variety of protein expression subpopulations of interest for diagnostics or therapeutics application.

## Materials and Methods

All chemical reagents and supplies were purchased from Millipore Sigma unless noted.

### EV Generation and Collection

The CELLine 1000AD bioreactor was utilized to maximize concentration of SK-OV-3 EVs and minimize processing steps as described previously^25,26^. SK-OV-3 cells at passage 6 were plated and expanded in flasks to reach the minimum of 2.5×10^7^ cells required to seed the bioreactor. McCoy’s 5A, 1X (Iwakata and Grad Mod.) with L-glutamine supplemented with 10% fetal bovine serum (FBS) and 1% penicillin/streptomycin were used for media during expansion. At passage 9, 2.7×10^7^ SK-OV-3 cells were seeded in the cell compartment of a CELLine 1000AD bioreactor in 15 mL of supplemented media. 1L of McCoy’s 5A, 1X (Iwakata and Grad Mod.) with L-glutamine supplemented with only 1% penicillin/streptomycin was added to the media compartment. After 24 hours to ensure full adhesion of cells to the matrix in the cell compartment, cell compartment media was replaced with EV-depleted media. EV-depleted media was prepared by centrifuging McCoy’s 5A, 1X (Iwakata and Grad Mod.) with L-glutamine supplemented with 10% FBS overnight at 100,000xg. 1% penicillin/streptomycin was then added, and the media was filtered. After seven days the cell compartment media was extracted to collect EVs and replaced with EV-depleted media, and the media in the media compartment was also replaced. This was repeated each week for a total of 6 weeks.

Placental mesenchymal stem cells (pMSCs) were isolated and expanded as previously described^27^. Briefly, pMSCs were isolated from de-identified second trimester human placenta from the chorionic callus tissue after explant culture. These cells were expanded in HyClone Medium high glucose supplemented with 5% fetal bovine serum, 20 ng/mL basic fibroblast growth factor, 20 ng/mL epidermal growth factor, and 1% penicillin/streptomycin and expanded at low passage numbers. 48-96 h prior to EV collection, cells were collected, washed with phosphate buffered saline (PBS) and reseeded with EV depleted media as prepared above. Conditioned media was collected from pMSCs at passage 4 from 20-35 T150 flasks.

Serum samples were provided by the UC Davis Comprehensive Cancer Center biorepository as deidentified remnants, obtained from patients who received a clinician-ordered CA-125 test. From each patient, ~2 mL of serum was obtained.

### EV isolation

#### Ultracentrifugation

To isolate EVs, we followed optimized ultracentrifugation isolation procedures recently reported^28,29^. Approximately 15 mL of cell culture supernatant or 1 mL of serum was centrifuged at 300xg for 10 min at 4°C to remove any cells and large debris. The EV containing supernatant was collected and centrifuged at 2,000xg for 15 min at 4°C to remove apoptotic bodies and debris. The supernatant was further subjected to 10,000xg for 30 min at 4°C to remove larger microvesicles. This supernatant was spun twice at 120,000xg for 70 min at 4°C, with the pellet resuspended in DI water in between spins. The final supernatant was discarded, and the pellet resuspended in 125 - 225 μL of MilliQ water or PBS. This solution was aliquoted and frozen at −80°C until used. Freeze-thaw cycles were minimized for downstream application. Centrifuge steps for SK-OV-3 samples were completed in an Optima LE-80L centrifuge with a SW-28 rotor. Slow speed centrifuge steps for serum samples were completed a Beckman Coulter Microfuge 20R centrifuge with an FA361.5 Biosafe rotor and slow speed spins were completed in a Beckman Optima TLX Ultracentrifuge with a TLA 100.1 fixed angle rotor.

pMSC EVs followed the same general protocol with two minor differences. First, after the 2,000xg spin two filtration steps were added. The supernatant was transferred to a 0.2 μm filter and vacuum filtered. The EVs in the filtrate were then concentrated in sterilized Centriprep 100,000 K cutoff filters at 8,836xg for 30 minutes, adding additional media until all media had been concentrated, before continuing on to the 10,000xg spin. Finally, the samples were subjected to 120,000xg for 90 minutes each instead of 70 minutes during the last two steps. Centrifuge steps for pMSC samples were completed in an L7 Ultracentrifuge with an SW-28 rotor (Beckman Coulter).

#### Size Exclusion Chromatography (SEC)

Prior to SEC, samples were centrifuged as described in the ultracentrifugation isolation methods through the 10,000xg spin. Izon 70 nm qEV_single_ columns were utilized in the Izon Automatic Fraction Collector (AFC). Columns were rinsed using 4 mL of 0.2 μm-filtered PBS. After rinsing, any remaining solution was removed from the top of column, 150 μL of sample was placed on top of the column and allowed to enter before adding filtered PBS. After void fraction elution, 2 fractions of 0.2 mL were collected that were confirmed to contain EVs in prior experiments. These fractions were combined and frozen at −80°C until used.

### EV characterization

#### Nanoparticle Tracking Analysis (NTA)

EVs were diluted in DI water to be analyzed by the NanoSight LM10 (Malvern Panalytical Ltd., UK)for size and concentration determination. The instrument was equipped with a 405 nm laser module and sCMOS camera. The NanoSight tubing was rinsed with filtered water prior to use. An accompanied automated syringe pump (Harvard Bioscience, MA, USA) allowed for recording a statistically representative portion of the EV isolate in flow conditions instead of recording the videos in a stagnant view. The particle concentrations were diluted to 1×10^8^ and 1.6×10^9^ part/mL in order to acquire an optimal readout for the data processing. The average concentration was also used for ExoView chip dosing. 3×30 second videos were obtained at camera level 10-11, and the data was analyzed using NanoSight NTA 3.1. software (detection threshold 2-3).

#### Transmission Electron Microscopy

Stock EVs were diluted 1:1 in 4% paraformaldehyde. A 10 μL drop of this solution was placed on the back side of parafilm covering a glass slide. A copper TEM grid was then floated on top of the EV solution for 20 min at room temperature. The grid was transferred to a 100 μL drop of TEM grade PBS to wash and then quickly to 2% glutaraldehyde for 5 min. The grid was then transferred to 8 successive 100 μL DI water drops for 2 min each to wash. The grid was then transferred to a 50 μL drop of uranyl oxalate for 5 min. Each grid was then dried by wicking the solution on filter paper. EVs were then imaged by transmission electron microscopy on a Talos L120C (Thermo Fisher Scientific, MA, USA) within 24 h.

#### Western Blot

EV pellets were lysed in RIPA lysis buffer containing protease inhibitor cocktail (Roche). A total of 2 μg of protein per lane was loaded on a 4-12% graded Tris-glycine SDS-polyacrylamide gel and proteins were transferred to a 0.2μm PVDF membrane (Life Technologies). Membranes were blocked in 5% milk in PBS with 0.05% Tween-20 for one hour at room temperature with rocking, and then incubated with primary antibody in blocking buffer at 4°C overnight. Antibodies to CD9 were used at 1:1000 dilution (BioLegend), CD63 at 1:1000 dilution (BD Biosciences), and CD81 (Santa Cruz Biotechnology) at 1:500 dilution. Secondary antibody mouse IgGκ-HRP was diluted at 1:5000 in block buffer and incubated at 1 hour at room temperature with rocking (Santa Cruz Biotechnology). Proteins were visualized with Supersignal West Pico PLUS chemiluminescent substrate (ThermoFisher) using a KwikQuant imager (Kindle Biosciences).

#### ExoView Tetraspanin Kit Assay

Tetraspanin Kits, for *in vitro* samples, and Plasma Kits, for clinical samples, were used as purchased (NanoView Biosciences) to interrogate EVs. All chips were stored at 4°C when not in use and allowed to warm to room temperature prior to use. Chips were pre-scanned using the provided protocol to identify any previously adhered particles during manufacturing. For incubation, chips were placed in wells of a 24 well plate, avoiding contact of the chip corners with the sides of the well. Water was added to the void space of the 24 well plate to dampen vibrations. EVs were diluted to the stated concentration, or at least 1:1, in the provided Incubation buffer and 35 μL of this solution carefully pipetted directly onto each chip, avoiding allowing the solution to spill off the edges. The plate was then covered in aluminum foil and allowed to incubate at room temperature overnight. The following morning, 1 mL of incubation solution was added to each chip and the plate was shaken at 500 rpm for 3 min. 750 μL of solution was removed from each well, replaced with 750 μL of new incubation solution, and shaken at 500 rpm for 3 min. This was repeated twice more for a total of 4 shake steps. During shake steps, detection antibody solution was prepared by combining incubation solution and blocking solution in a 2:1 ratio and adding each of the provided tetraspanin antibodies in a ratio of 1 μL:600 μL solution. The provided tetraspanin panel includes: CF488-anti-CD9, CF647-anti-CD63, and CF555-anti-CD81. After the last shake, 750 μL of solution was removed and 250 μL of antibody solution was added to each well. The plate was covered in aluminum foil and allowed to incubate at room temperature for 1 h. After incubation, 500 μL of incubation solution was added to each well. Immediately following, 750 μL of solution was removed from each well and replaced with 750 μL of new incubation solution. The plate was then shaken at 500 rpm for 3 min. 750 μL was removed from each well, replaced with 750 μL of wash solution, and shaken at 500 rpm for 3 min. This was repeated 3 times for a total of 3 shakes after addition of wash solution. 750 μL was removed from each well, replaced with 750 μL of rinse solution, and shaken at 500 rpm for 3 min. Two petri-dishes were filled with rinse solution and each chip was transferred with flat tip tweezers from the 24 well plate to the first dish, swirled, then to the second dish and swirled. During each transfer, care was taken to ensure the chip remained flat so that the antibody array remained wet. Chips were dried by tipping the chip at a 45° angle, slowly lifting out of solution, and placing on Kim wipe. The chips were then transferred to the chuck and scanned for interferometric and fluorescence imaging. During data analysis, fluorescence cut-offs were chosen by limiting the number of detected particles on MIgG capture spots to approximately 10 events. For the tetraspanin antibodies, we used fluorescence cutoffs of 400 a.u. for the red and green channels and 600 a.u. for the blue channel in all experiments. We have adopted any alterations to this protocol over time recommended by ExoView, checking that the apparent tetraspanin profile of SK-OV-3 EVs remains consistent after each update.

For ovarian cancer marker staining, AlexaFluor 488 anti-human CD326 (EpCAM), PE anti-human CD24, and AlexaFluor 647 anti-human CD340 (HER-2) were incubated at 1 μg/mL in blocking solution. Fluorescence cut-offs of 600 a.u., 300 a.u., and 300 a.u. were used for the blue, green, and red channels, respectively. This panel of antibodies was purchased from BioLegend.

#### Flow Cytometry

A CytoFLEX flow cytometer (Beckman Coulter) equipped with four lasers was used for all flow cytometry experiments. Prior to EV analysis, we employed a calibration procedure utilizing standard beads and FCM_PASS_ software described recently by Welsh et al^22^. For size calibration, we utilized a mixture of NIST Traceable polystyrene beads (ThermoFisher) including 100 nm, 152 nm, 203 nm, 269 nm, 345 nm, 401 nm, and 453 nm. 405 nm SSC was used for triggering and the threshold was set to achieve an event rate of approximately ~2000 events/s on the lowest flow rate of 10 μL/min when analyzing 0.2μm-filtered ultra-pure water. In this case the threshold was 1000 (a.u) at a gain of 200. Beads were diluted in 0.2 μm-filtered water to reach a total event rate of approximately 10,000 events/s and recorded for 120 seconds. These data were analyzed in FCM_PASS_ software to convert the arbitrary scatter intensity to nominal diameter of particles^30^. Gain of the various channels was optimized by analyzing 8 Peak Rainbow Calibration Beads from Spherotech (Cat. RCP-30-5A). Gain was increased incrementally from 25 to 3000 in each channel. The gain that best separated the dimmest bead population from background, calculated by stain index ((MFI_beads_-MFI_background_)/Stdev_background_), was chosen for further studies. Next, a combination of 200 nm and 500 nm fluorescent beads were utilized to optimize 405 nm SSC gain. This is a slight deviation from the procedure of Welsh et al as they utilized 100 and 200 nm beads^22^. Here, 405 nm SSC-H was used to trigger with a threshold of 900. Gain of 405 nm SSC was increased incrementally from 1 to 300 and the value that maximized the separation of the background 405 nm SSC intensity from the 200 nm beads was chosen, with a secondary goal of maximizing separation of 405 nm SSC intensity of 200 nm and 500 nm beads.

For antibody titrations, 1.0×10^9^ SK-OV-3 EVs were incubated with 0.5 μL – 8 μL of each antibody provided by NanoView, described in ExoView Tetraspanin Kit Assay methods, and diluted to a total volume of 50 μL. Each sample was allowed to incubate for 30 min at 4°C before dilution up to 2 mL in 0.2 μm-filtered PBS. Optics parameters for particle detection and thresholding were determined from the calibration and optimization as described above. Here we used 405 nm SSC to trigger with a threshold of 900 with a gain of 100 and the following fluorescent channel gains: 488 nm laser: 525/40 nm filter gain of 2000, 561 nm laser: 585/42 nm filter gain of 1500, 638 nm laser: 660/20 nm filter gain of 1000. This produced approximately 1000 events/s of noise and impurities when analyzing 0.2 μm-filtered PBS. EVs were analyzed at the lowest flow rate of 10 μL/min for 3 min. 1 μL, 2 μL, and 3.5 μL of anti-CD9, anti-CD63, and anti-CD81 antibody, respectively, was found to be the best volume to maximize separation index. Using this volume, either 1.0×10^9^ of SK-OV-3, pMSC, or serum EVs were incubated and analyzed in the same conditions described previously. In addition, a sample of 1.0×10^10^ serum EVs was incubated with antibodies, since lipoprotein co-isolated with EVs and can dramatically inflate the concentration when analyzed by NTA. In addition to the typical dilution to 2 mL, the SK-OV-3 EV sample was also analyzed at dilutions of 0.5 mL, 1 mL, 4 mL, and 8 mL to check for swarming, the interrogation of multiple EVs at a single time.

#### Biotinylation of EVs

Biotin was covalently coupled to free amines of exposed EV membrane proteins in a non-specific manner. To accomplish this, 10^9^ - 10^10^ EVs (as assessed by NTA) were incubated with 0.5 mM EZ-Link™ Sulfo-NHS-LC-Biotin in 50 μL of PBS at room temperature for 30 min. Biotinylated EVs were diluted to 150 μL and separated from free biotin by SEC using qEV_single_ columns on the Izon AFC. The first 0.2 mL fraction was diluted 1:20 with Solution A and incubated on chips as described.

#### Statistical methods

All experiments were performed in triplicate and error bars represent one standard deviation (s.d.) from the mean. Statistical significance was assessed using built-in functions of MATLAB 2020a Update 5 software. For comparison of two populations, a Student’s t-test was used, while ANOVA was used for three or more populations.

## Results

### EV Generation, Isolation, and Characterization

We first investigated the consistency of EV tetraspanin expression across biofluid source and isolation method. As sources, we chose isolated EVs from pMSCs, due to their known therapeutic applications, as well as ovarian cancer (OvCa) cells and serum from OvCa human patients, due to their utility for clinical diagnostics. For isolation methodology, we chose either ultracentrifugation (UC), as previously described^29^, or size exclusion chromatography (SEC) since these are common isolation methods used by EV researchers^31^. UC and SEC are known to enrich small EVs, which contains the canonical exosomes, a subclass of EVs defined by their biogenesis via inward budding into multivesicular bodies (MVBs) and subsequent release to extracellular space. However, since other types of EVs, such as microvesicles, ectosomes, exomeres, etc., have overlapping size and density, and there are no known exosome-exclusive markers or features, it is not currently possible to ensure that all isolated EVs originate from the MVBs. Therefore, in this work, we more generally apply the term “EVs” to encompass the diverse array of particles isolated by either UC or SEC.

EVs produced from SK-OV-3 cells, a human OvCa cell-line, were utilized to first optimize our incubation procedure for immunocapture. SK-OV-3 cells were grown in a bioreactor, which allows for continuous collection of EVs at high concentrations^25,26^. EVs were isolated from the conditioned media in the cell compartment each week^25^ and extensively characterized to confirm the presence of EVs. TEM images revealed particles of varying size with a deflated cup shape, an artefactual morphology characteristic of EVs when imaged by TEM (**Figure 1a)** ^32,33^. In accordance with the *Minimum Information for Studies of Extracellular Vesicles 2018* (MISEV2018) guidelines^10^, the presence of proteins was confirmed by Western blot for CD9, CD63, and CD81 (**Figure 1b**), and NTA confirmed that EVs were isolated at a high concentration with a peak diameter of 128.8 nm (**Figure 1c**).

**Figure 1:**
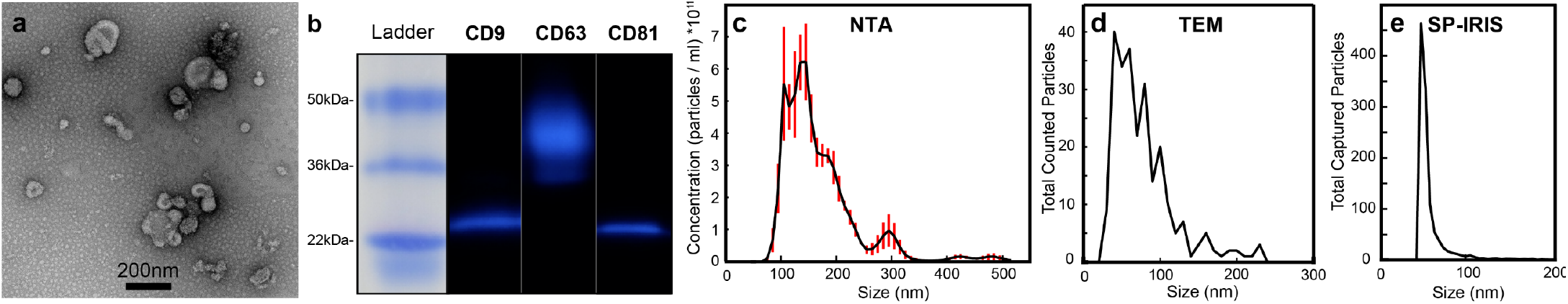
Characterization of SK-OV-3 EVs isolated from bioreactor conditioned media. (a) TEM micrograph of isolated EVs. (b) Western blot of EV associated tetraspanins CD9, CD63, and CD81. Size distribution was assessed by either (c) NTA, (d) TEM, or (e) SP-IRIS (anti-CD81 capture). The distributions, modes, and apparent concentrations varied between the three techniques. Error bars represent +/- one s.d.

We compared the apparent size distributions of SK-OV-3 EVs between NTA, TEM, and SP-IRIS. TEM (**Figure 1a,d**, with additional micrographs in **Supplementary Figure 1**) is the closest to ground truth, with a mode diameter of ~35 nm and 50% of total EVs by count in the size range from 40-90 nm. NTA exhibits known artefacts in measuring EV size, given that it is only capable of detecting EVs larger than ~70 nm in nominal diameter due to the low scattering of organic particles^15^. This results in an apparent mode between 100-150 nm (**Figure 1c**), not in agreement with TEM measurement, which shows a log-linear particle count vs. size down to ~30 nm (**Figure 1d**)^15^. SP-IRIS size distributions matched well with our TEM results, showing a mode diameter of 50 nm (representative size data for anti-CD81 capture is shown) and a similar size profile to TEM, i.e., a logarithmic-linear decrease in count over the diameter range of 50-100 nm (**Figure 1e**).

### Optimizing incubation concentration to identify single-EVs

In SP-IRIS, EVs are incubated on a microarray of antibody spots against CD9, CD63, CD81, or mouse immunoglobulin G (MIgG, a non-specific binding control). Captured particles can be measured in two ways: (1) count and multiplexed marker expression by immunofluorescence (**Figure 2a**), or (2) count and size by interferometry (SP-IRIS, **Figure 2b**). For fluorescence detection, the ExoView R100 permits detection of up to three color channels following immunocapture. Fluorescence is captured following excitation by LED and images are superimposed to identify co-expression of each protein on a single EV (**Figure 2c**).

**Figure 2:**
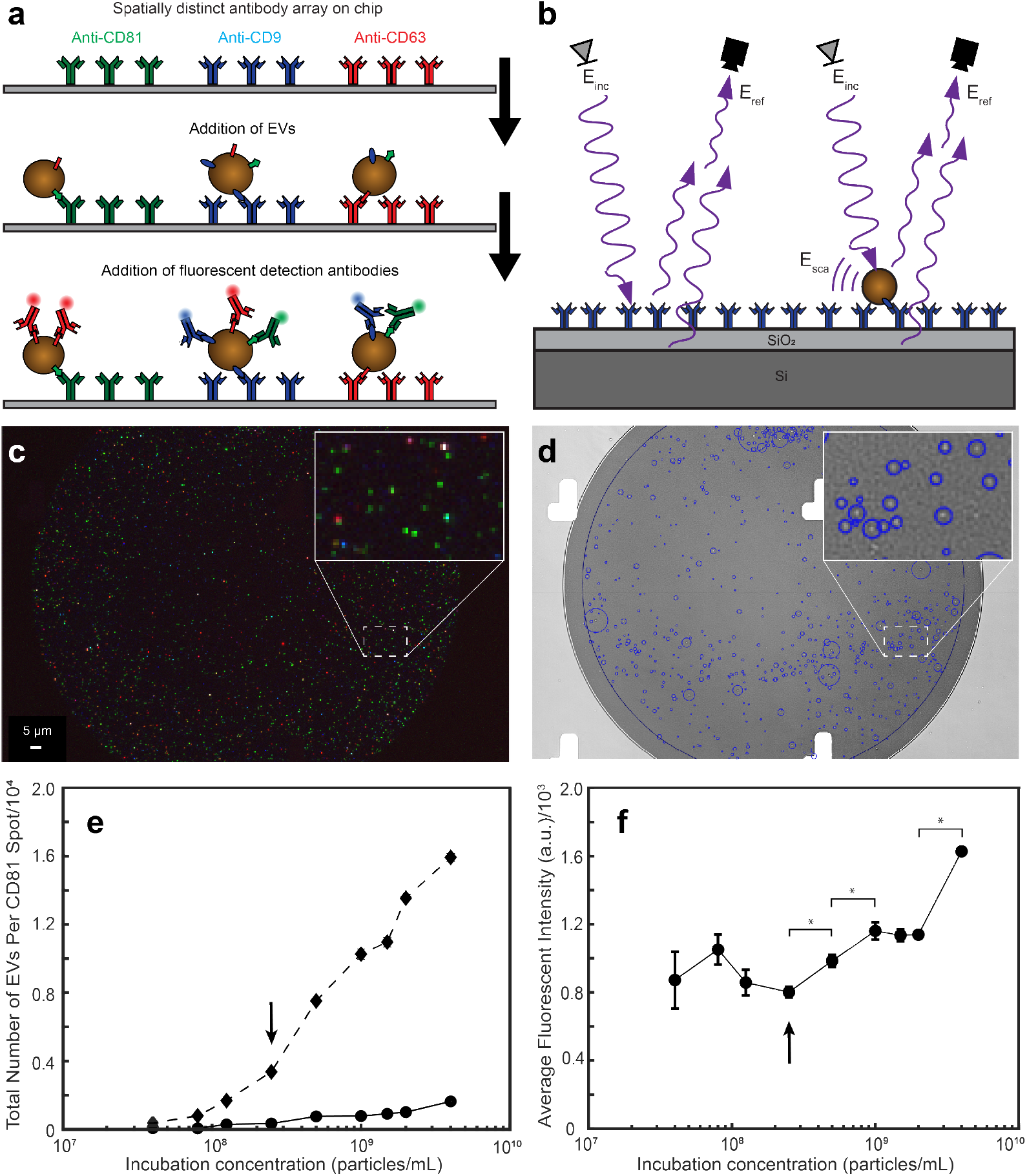
ExoView platform for particle capture and analysis. Immuno-captured EVs are characterized by (a) immunofluorescence and (b) SP-IRIS. Images captured in either (c) fluorescence mode or (d) SP-IRIS mode can indicate detected particles (blue circle diameter scaled to estimated EV diameter). (e) Serially diluted EVs were incubated on chips, with captured EVs detected by SP-IRIS (circles) or immunofluorescence (diamonds). Comparing counts between the two cases revealed a discrepancy in limit of detection, with fluorescence yielding higher sensitivity. (f) To find the optimal incubation concentration where signal was maximized for single particles, fluorescence intensity per particle was calculated. Arrows indicate the highest concentration with reliable single-particle detection, above which “swarming” occurs. Error bars represent +/- one s.d. * p<0.05 by Student’s t-test.

SP-IRIS mode uses the interference of two reflected light paths illuminated by LED, one that passes through a bound particle and another through an empty part of the thin film silicon chip (**Figure 2b**)^24,23^. The resulting interference pattern provides increased resolution beyond the diffraction limit, and in principle does not have a particle size limitation^24^. Due to potential issues with sample drift over long integration times required for single particle imaging, the ExoView platform currently features a practical detection limit down to ~50 nm, assuming a constant refractive index of 1.4 (**Figure 2d**).

Although some reports of EV analysis using the new ExoView platform are beginning to emerge^24^, to our knowledge, there is no reporting of optimization for reliable detection of single particles. Therefore, prior to analyzing the tetraspanin profile of EVs, we identified the optimal concentration of SK-OV-3 EVs that maximized EV count while ensuring that each identified particle was a single EV.

Using a standard tetraspanin microarray featuring four repeated spots each of anti-CD63, anti-CD9, anti-CD81, and anti-MIgG, we performed a serial dilution on SK-OV-3 EVs, ranging from bulk concentration of 4×10^7^ to 4×10^9^ particles/mL (as measured by NTA). First the number of EVs detected by each method was compared to identify the sensitivity of each method. Number of EVs identified by fluorescence was determined by adding all particles with higher fluorescent intensity than the MIgG spot in one or more channels. In interferometry-based SP-IRIS mode, approximately 10x fewer EVs were detected compared to fluorescent detection (**Figure 2e**). As the largest number of SK-OV-3 EVs were identified by anti-CD81 capture/CD81 detection, this combination was used for this analysis. Above a concentration of 2.5×10^8^ particles/mL, the particle counts no longer doubled with a doubled incubation concentration; therefore, it was not clear if measured spots were truly single EVs or fluorescence from multiple particles, also known as “swarming.”

To identify whether this apparent upper concentration limit was due to binding limitations (e.g., steric hindrance or lack of available binding sites) or instead due to fluorescence overlap, the average fluorescent intensity of the particles at each concentration was identified. If measured spots are truly single particles, as incubation concentration increases, particle count should increase, but the fluorescent intensity of each particle should remain the same. Increasing incubation concentration incrementally from 4.0×10^7^ to 2.5×10^8^ particles/mL produced no significant increase in fluorescent intensity per bound particle. However, above incubation concentrations of 2.5×10^8^ particles/mL, there was a significant increase in fluorescent intensity per bound particle (**Figure 2f**). From this point forward, we used 2.5×10^8^ particles/mL as the optimal incubation concentration for EVs. Practically, this translated to ~2500 captured particles/spot.

### SK-OV-3 EVs consistently exhibit time-independent tetraspanin expression

Next we analyzed tetraspanin profiles for SK-OV-3 EVs isolated by UC. SK-OV-3 EVs expressed all three tetraspanins with varying co-expression of each. We measured discrepancies between capture and detection, even for the same tetraspanin. For example, the highest number of detectable EVs were captured by anti-CD9, although very few were detected by anti-CD9 on any capture spot (**Figure 3a,** blue bars). On the other hand, the majority of EVs captured across each of the tetraspanins were detected by anti-CD81 in all cases (**Figure 3a**, green bars). Compared to anti-CD81 capture, similar numbers of EVs were captured by anti-CD63, yet notably fewer anti-CD63-captured EVs were detected by anti-CD9.

**Figure 3:**
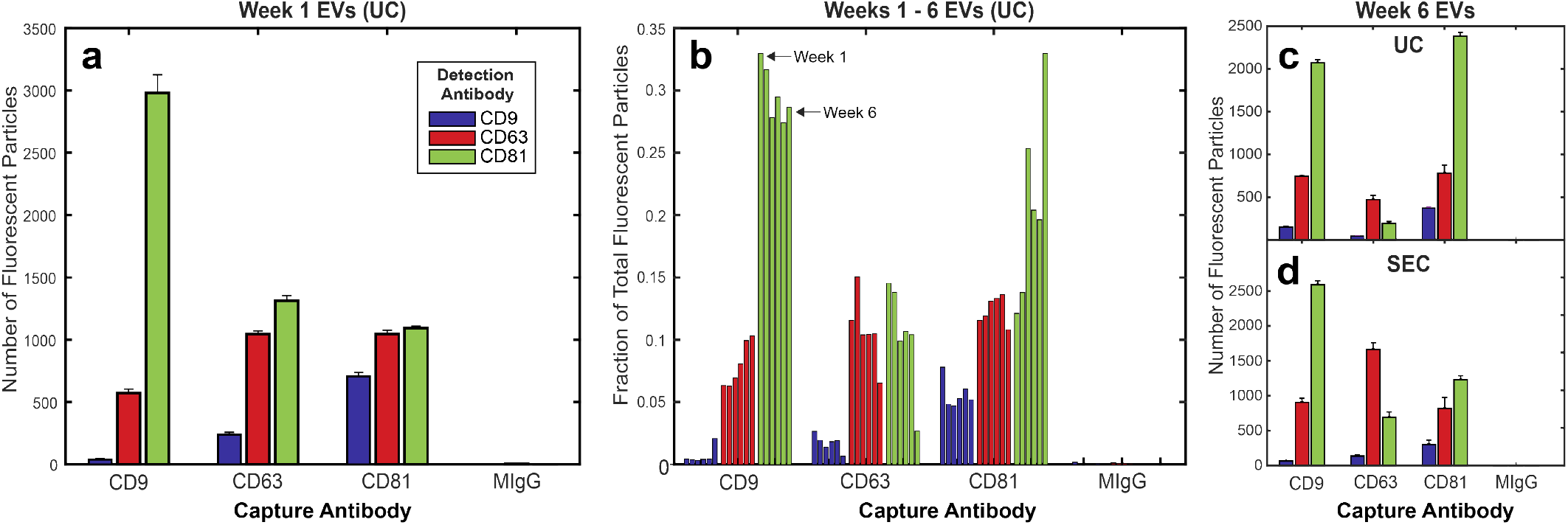
Fluorescence profiling of tetraspanin expression for isolated SK-OV-3 EVs. (a) Representative example of tetraspanin colocalization for SK-OV-3 EVs isolated by UC after 1 week of cell growth in a bioreactor flask. (b) Tetraspanin colocalization pattern of SK-OV-3 EVs remained relatively consistent across 6 consecutive weeks of isolation by UC from bioreactor grown cells. Tetraspanin colocalization pattern of SK-OV-3 EVs from week 6 of the bioreactor isolated by (c) UC or (d) SEC showed some differences but did not have variation outside that seen from week to week of the bioreactor isolations. Error bars represent +/- one s.d.

Next, we measured whether this profile was consistent amongst EVs isolated from cells grown in a bioreactor flask over the course of several weeks (**Figure 3b**). The bioreactor allows for collection of EVs in volumes of ~15 mL from over 2.5×10^7^ cells in 3D culture with minimal upkeep, and has been adopted by multiple labs ^25,34^. Since this or similar methods will likely be utilized for scale-up in industrial application, it is important to identify the consistency of EVs from this type of system from consecutive harvests. Although EVs from each week were incubated at 2.5×10^8^ particles/mL on each chip, the total number of fluorescently identified EVs varied by week. For example, CD81 capture/CD81 detection ranged from 1000 – 4500 total particles depending on the week. This may have been due to low precision associated with NTA derived concentrations used to inform chip incubation concentration. For this reason, we normalized the tetraspanin profiles of the EVs from each week to the total number of EVs fluorescently detected per chip (**Figure 3b**). Therefore, when normalized to be independent of total EV yield over the course of 6 weeks, SK-OV-3 EVs showed relatively low variability in tetraspanin profile, with less than a 5% standard deviation for most capture/detection combinations. One exception was detection of anti-CD81 for particles captured by anti-CD81, with a slightly higher variation of 7.68% across 6 weeks. EVs produced from SK-OV-3 across different bioreactors from various cell passages were tested and showed similar tetraspanin profiles (**Supplementary Figure 2**).

### Effect of isolation method and biofluid source on tetraspanin profile

Given the reported effects on EV composition according to isolation methodology^35,36^, we measured EV tetraspanin profile using SEC or UC. After UC, SEC is the next most common method to isolate EVs according to a recent poll^31^. To directly compare UC to SEC, conditioned media from the bioreactor containing SK-OV-3 cells at week 6 was split in half and isolated either by UC or SEC. EVs isolated from either method exhibited a similar tetraspanin profile i.e., for each capture spot, the particle count of each tetraspanin followed the same relative trend (**Figure 3c,d**). Notably, total counts by anti-CD63 capture increased for SEC isolated EVs, while the number of anti-CD81 captured particles decreased. However, these differences were not outside of the week-to-week variation of bioreactor samples isolated by UC (**Figure 3b**).

After identifying that SK-OV-3 EVs had a consistent tetraspanin profile across different isolation weeks and different isolation techniques, we expanded to other biofluid sources to find if this trend is indicative of all EVs. We isolated EVs from pMSCs, chosen for clinical applicability. MSC-derived EVs have a variety of therapeutic effects and have been of great interest for treating numerous diseases. pMSC EVs in particular have neuroprotective properties and are being studied for treatment of spina bifida and multiple sclerosis^37,38^. We also isolated EVs from human serum, to represent a more complex source with relevance to diagnostic analysis.

Each EV source has heterogeneous apparent tetraspanin profiles depending on capture antibody (**Figure 4a-c**). Detection of EVs by fluorescent antibody was compared both by the total number of fluorescent particles on any capture spot (**Figure 4d**) or separated by specific capture spot (**Figure 4e**). While some differences in anti-CD9 and anti-CD81 detection were apparent independent of capture spot, there was no significant difference in CD63 detection (**Figure 4d**). However, when separated by capture spot, there are distinct differences in multiplexed tetraspanin expression, suggesting that distinct subpopulations exist that are unique to each source (**Figure 4e**). Serum EVs exhibited relatively high anti-CD9 detection (**Figure 4d**), with a particularly distinct anti-CD9 capture/anti-CD9 detection peak (**Figure 4e**). This suggested that serum EVs exhibit a higher density of co-expressed CD9, since EVs were detected in higher numbers by anti-CD9 compared to SK-OV-3 EVs (**Figure 4a,b**). *In vitro* isolated pMSC EVs had a tetraspanin profile more similar to SK-OV-3 EVs with similar distinct CD9 capture and CD9/CD63/CD81 detection profiles (**Figure 4e**). However, pMSC EVs had a higher percentage of EVs that were detected by anti-CD9 (**Figure 4d**), suggesting that CD9 was more densely expressed on pMSC EVs compared to SK-OV-3 EVs, which was mainly due to CD81 captured EVs (**Figure 4e**).

**Figure 4:**
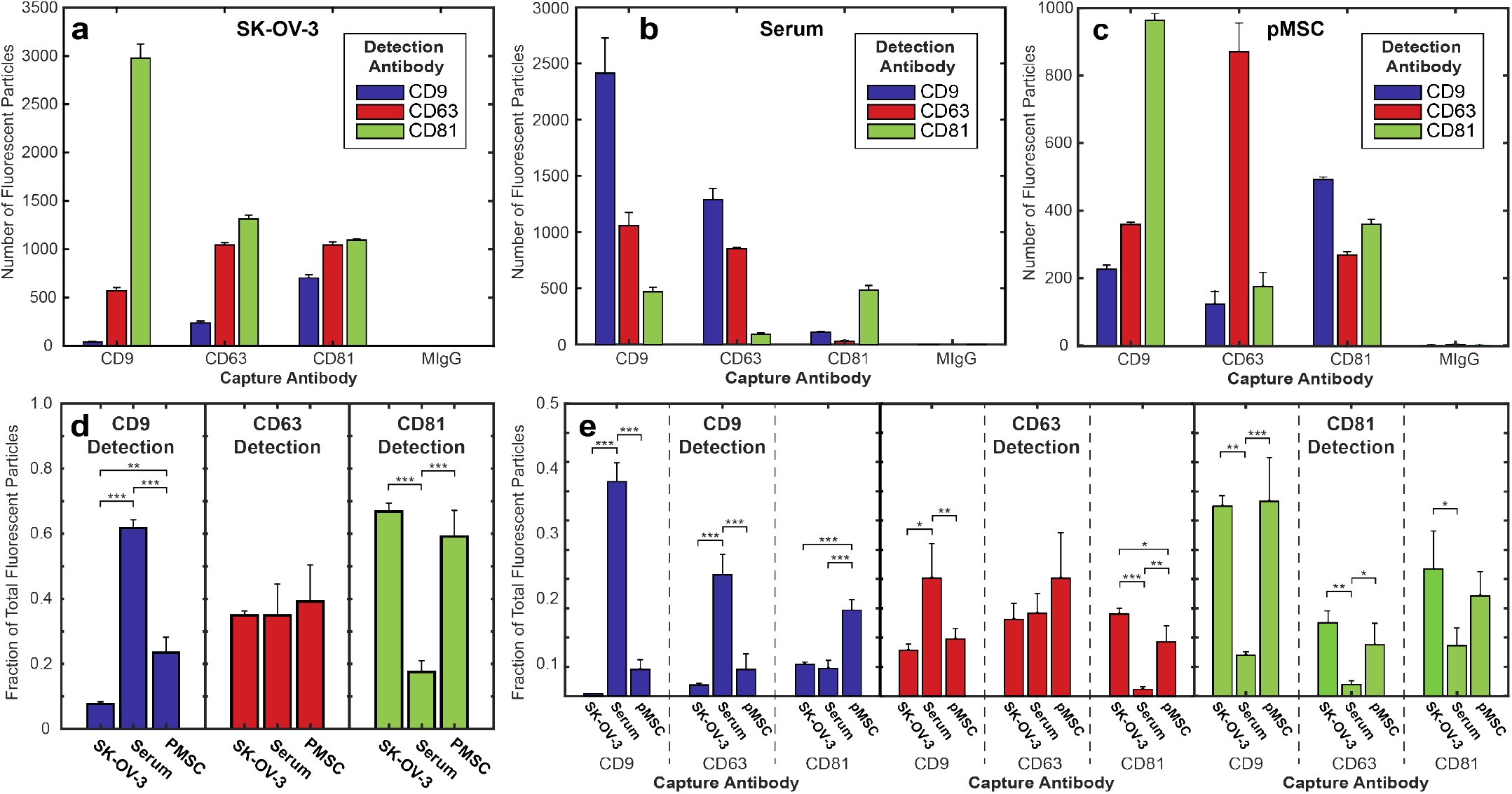
Comparison of tetraspanin profile of EVs isolated by UC from varying biofluid source. Representative EVs derived from (a) SK-OV-3, (b) serum, and (c) pMSCs exhibit distinct tetraspanin profiles. Three samples of EVs from each source were analyzed and the average fraction of total fluorescent particles was determined for detection antibody on (d) any capture spot or (e) single capture spot. This highlights the complex nature of tetraspanin multiplexing. Error bars represent +/- one s.d. * p<0.05, ** p<0.01, *** p<0.001 by ANOVA.

### Single particle flow cytometry to complement tetraspanin expression

Nanoscale flow cytometry is a complementary technique to analyze single-EV protein expression. In comparison to fluorescence microscopy of EVs by ExoView, flow cytometry is much higher throughput, allowing interrogation of ~100,000 particles/min with a variety of options for fluorescence detection (more light sources/filters). The main drawback of flow cytometry is that it is scatter limited, producing a theoretical limit of detection of ~80 nm for EVs in the best case. We directly compared tetraspanin expression of the aforementioned various EV populations between SP-IRIS and nanoscale flow cytometry to compare apparent tetraspanin profiles.

Prior to analysis of EVs, we calibrated the flow cytometer using a variety of standard refractive index, size, and fluorescent beads, as described by Welsh et al^30^. This allows us to produce standardized, calibrated data and to optimize the instrument for detecting nanoparticles^22,39^. To better visualize the limit of detection, the 405 nm SSC-H threshold was chosen to visualize approximately 1000 events/s of electronic noise (**Figure 5a**). This procedure permits tuning of the detection threshold above the electronic noise to maximize the number of detected EVs. For light scatter calibration, we analyzed the scattering intensity of a variety of polystyrene beads. These scattering intensities of beads of known diameter could then be used to estimate the size of particles of constant refractive index based on Mie scattering theory using publicly available FCM_PASS_ software. Using this software and optimized gains, we determined our instrument’s theoretical detection limit by transforming the arbitrary units of SSC intensity to units of size, assuming a constant refractive index (details shown in **Supplementary Figure 3**)^30^. With a threshold of 900 for triggering by SSC-A, we found that our reliable limit of detection when gating for EVs above instrument noise was ~100 nm (**Figure 5b**).

**Figure 5:**
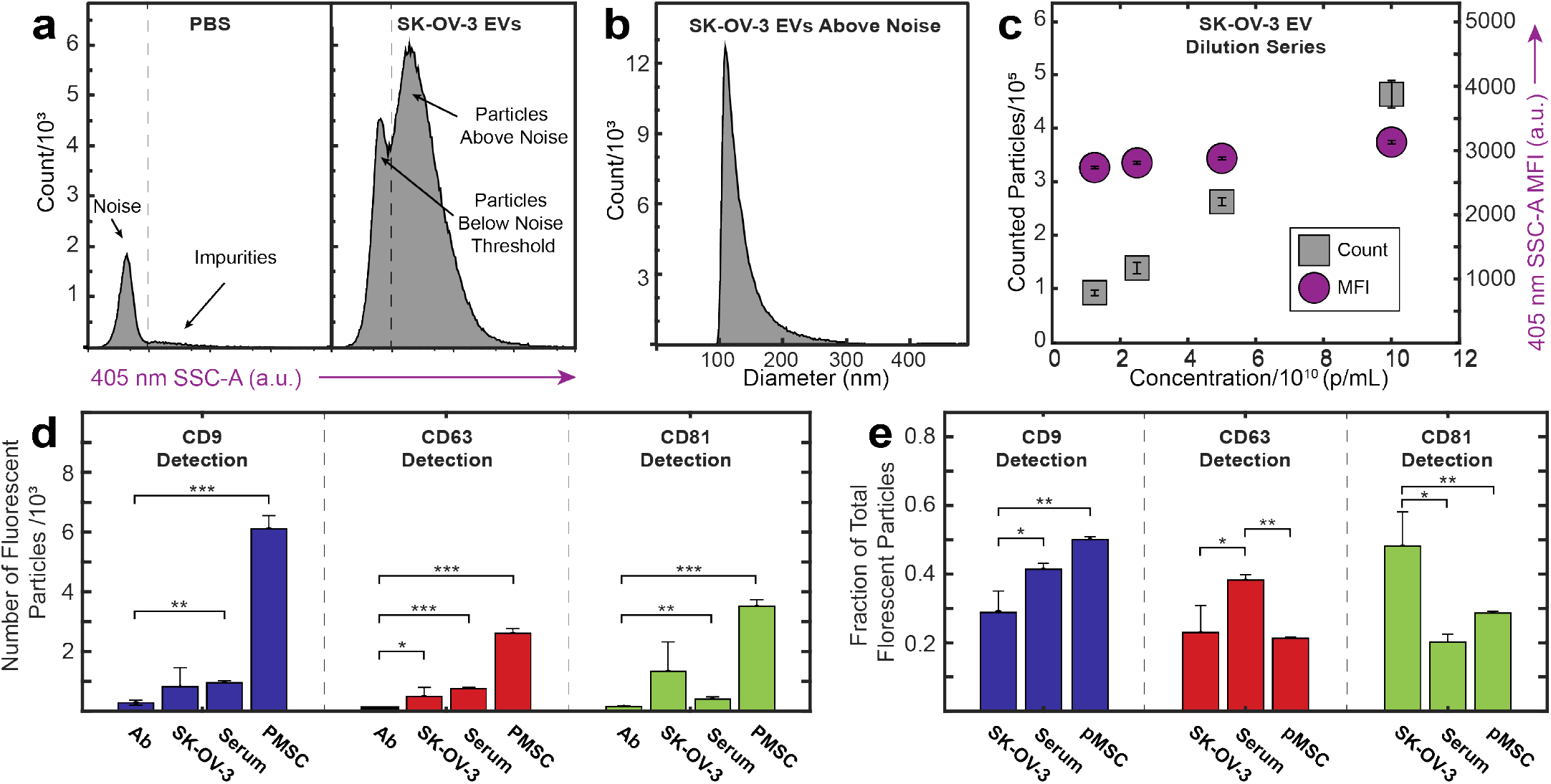
(a) Histograms of side scatter area (SSC-A) of control PBS, showing the gating choice above electronic noise and below impurities, and SK-OV-3 EVs, showing the additional events both within and above the noise limit. (c) Histogram of SK-OV-3 EV size converted to diameter using FCM_PASS_ calibration. (d) Dilution series of SK-OV-3 EVs illustrating constant median fluorescent intensity (MFI, violet circles) as events increased with concentration (gray squares), confirming that detected events are single particles in the concentration range tested. (d) Total number of fluorescent particles above background for labeled EVs versus antibody control. (e) Percent of total fluorescent EVs identified by each tetraspanin. Error bars represent +/- one s.d. * p<0.05, ** p<0.01, *** p<0.001 by Student’s t-test versus Ab control for (d) and ANOVA for (e).

To ensure that events were single EVs and not “swarming” EVs^40^ (multiple EVs being interrogated at one time producing a single event) labeled SK-OV-3 EVs were diluted in series. At all concentrations tested, total events decreased linearly with dilution, while scatter intensity remained constant, suggesting that at these concentrations, events represented single EVs (**Figure 5c**).

SK-OV-3, pMSC, and serum-derived EVs were analyzed for tetraspanin expression using the same fluorescently-labeled detection antibodies for the preceding ExoView experiments, to minimize any variability as a result of batch-to-batch differences. A final concentration of 5×10^8^ particles/mL, measured by NTA, was utilized for SK-OV-3 and pMSC EVs. Serum EVs were analyzed at 5×10^9^ particles/mL to achieve the same counts, likely due to co-isolated non-EV particles (e.g., lipoprotein particles) that are indiscernible from EVs by NTA. Gating schemes were chosen above background noise (**Supplementary Figures 4** and **5**). For SK-OV-3, pMSC, and serum-derived EVs, respectively, 19,600, 19,100 and 64,900 events, representing 2.5%, 1.3%, and 13.6%, of all events above background SSC intensity were above background fluorescence intensity in one or more channels.

Student’s t-tests were applied to assess statistical significance of detected events compared to the background noise events. For all tetraspanins, pMSC EVs had a significantly higher percentage of EVs above background (**Figure 5d**). Both SK-OV-3 and serum-derived EVs had populations of EVs that were discernable above non-labeled EVs and antibody only controls, although close to the limit of detection. Notably, SK-OV-3 CD9+ and CD81+ EVs were not statistically significant to the number identified in the antibody only control mainly due to low counts in one of the three replicates. The low amount of highly fluorescent CD9+ EVs was similar to the trends seen with SP-IRIS. Similar to our approach to display this complex multiplexing data in **Figure 4e** above, **Figure 5e** shows a complementary plot of the flow cytometry data with counts normalized to total particles, in order to visualize relative trends of detected particles. Serum EVs showed similar protein expression by flow cytometry and immunocapture, with the smallest fraction of EVs being CD81+. However, flow cytometry did not produce an apparent profile of pMSC EVs that was similar to immunocapture. For flow cytometry, although there were more CD81+ EVs compared to CD63+ EVs, the largest fraction was CD9+. By immunocapture, the fewest number of EVs were detected by CD9 fluorescence.

### Non-specific EV capture and profiling by EV biotinylation

For serum and pMSC derived EVs, EVs captured by different tetraspanins had distinct apparent fluorescent tetraspanin profiles. From this, we hypothesized that non-specific capture would provide a more encompassing view of the tetraspanin profile of the entire population. We realized non-specific capture of EVs via covalent biotinylation of exposed amine groups on EVs’ surface protein^41^. Customized chips were obtained from NanoView with an anti-biotin coating to facilitate non-specific capture. Biotinylated-EVs were separated from unbound biotin by SEC. Purified, biotinylated EVs were incubated on the anti-biotin coated chip and probed with the same tetraspanin detection panel used throughout the study, anti-CD9, anti-CD63, and anti-CD81 (**Figure 6a**). More EVs were detected on anti-biotin capture spots compared to anti-CD81 capture spots, which had the most numerous captures of any tetraspanin spot. These appeared to be true biotinylated EVs, as nonbiotinylated EVs nor the biotinylation agent itself identified any fluorescent particles. Furthermore, the biotinylation did not appear to significantly impact the apparent tetraspanin profile in most cases when compared to a non-biotinylated sample (e.g., **Figure 6a** “EV” vs “B-EV”).

**Figure 6:**
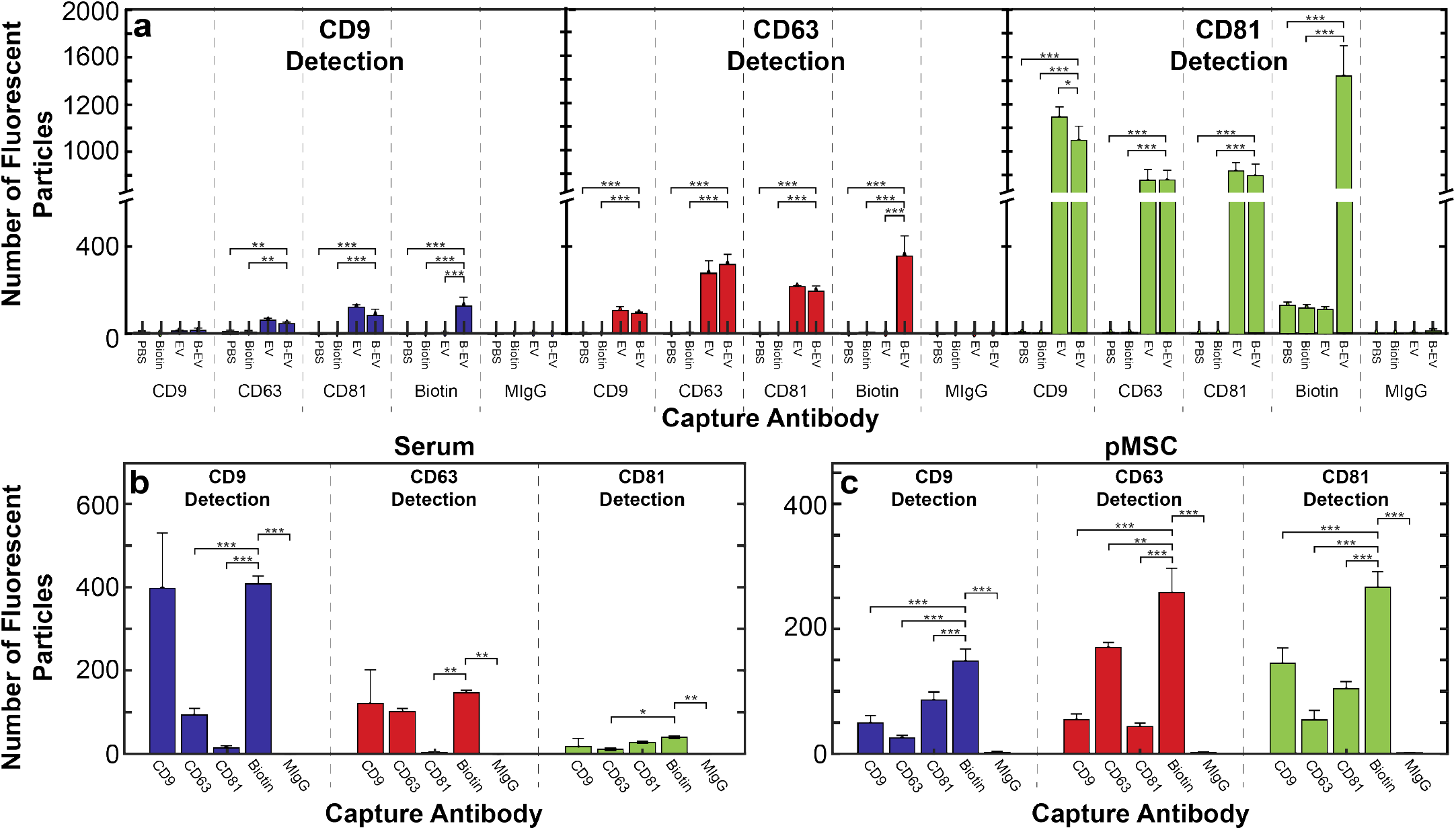
Biotinylated EV captured by tetraspanins and anti-biotin. (a) Biotinylated SK-OV-3 EVs (“B-EV”) were compared to PBS, free biotinylation agent (“Biotin”), and non-biotinylated SK-OV-3 EVs (“EV”). These results indicate that biotinylation did not majorly alter tetraspanin capture (i.e., “B-EV” compared to “EV”) in most cases. For biotinylated (b) serum and (c) pMSC derived EVs, some “multiplex bias” could be observed, e.g., CD9 captured by single-tetraspanin compared to non-specific capture. Error bars represent +/- one s.d. * p<0.05, ** p<0.01, *** p<0.001 by ANOVA.

Serum and pMSC derived EVs were also biotinylated and analyzed. Serum EVs, unlike SK-OV-3 EVs, did not show an overall increase in capture by anti-biotin compared to each single tetraspanin, but were captured as efficiently by anti-CD9 as by anti-biotin (**Figure 6b**). Furthermore, the apparent tetraspanin profile appeared similar to that of the CD9 capture spot. pMSC EVs had a higher number of particles captured by anti-biotin than any single tetraspanin (**Figure 6c**). However, the apparent tetraspanin profile did not match any of the single tetraspanins, having a similar number of EVs detected by CD63 and CD81, and a lower number detected by CD9.

### OvCa markers on EVs captured non-specifically or by single tetraspanins

Circulating EVs are attractive next generation diagnostic biomarkers due to the fact that their biochemical signatures are indicative of their origin (are subsets of their parent cell) and disease state. EVs from epithelial OvCa are on the leading front of this research, and often are used as a model to show proof-of-concept for EV biosensing platforms. Yet there is little known about the heterogeneity of the diagnostically relevant markers on these EVs. For this reason, we wanted to identify if multiplex bias by tetraspanin capture could affect the identification of diagnostically relevant proteins on SK-OV-3 EVs. Three markers frequently associated with OvCa, CD24, EpCAM, and Her2^42^, have also been utilized to identify the diagnostic efficacy of OvCa EVs^43,44^.

SK-OV-3 EVs were biotinylated and incubated with chips as previously described, and then labeled with fluorophore-conjugated anti-CD24, anti-EpCAM, and anti-Her2 detection antibodies (**Figure 7**).

**Figure 7:**
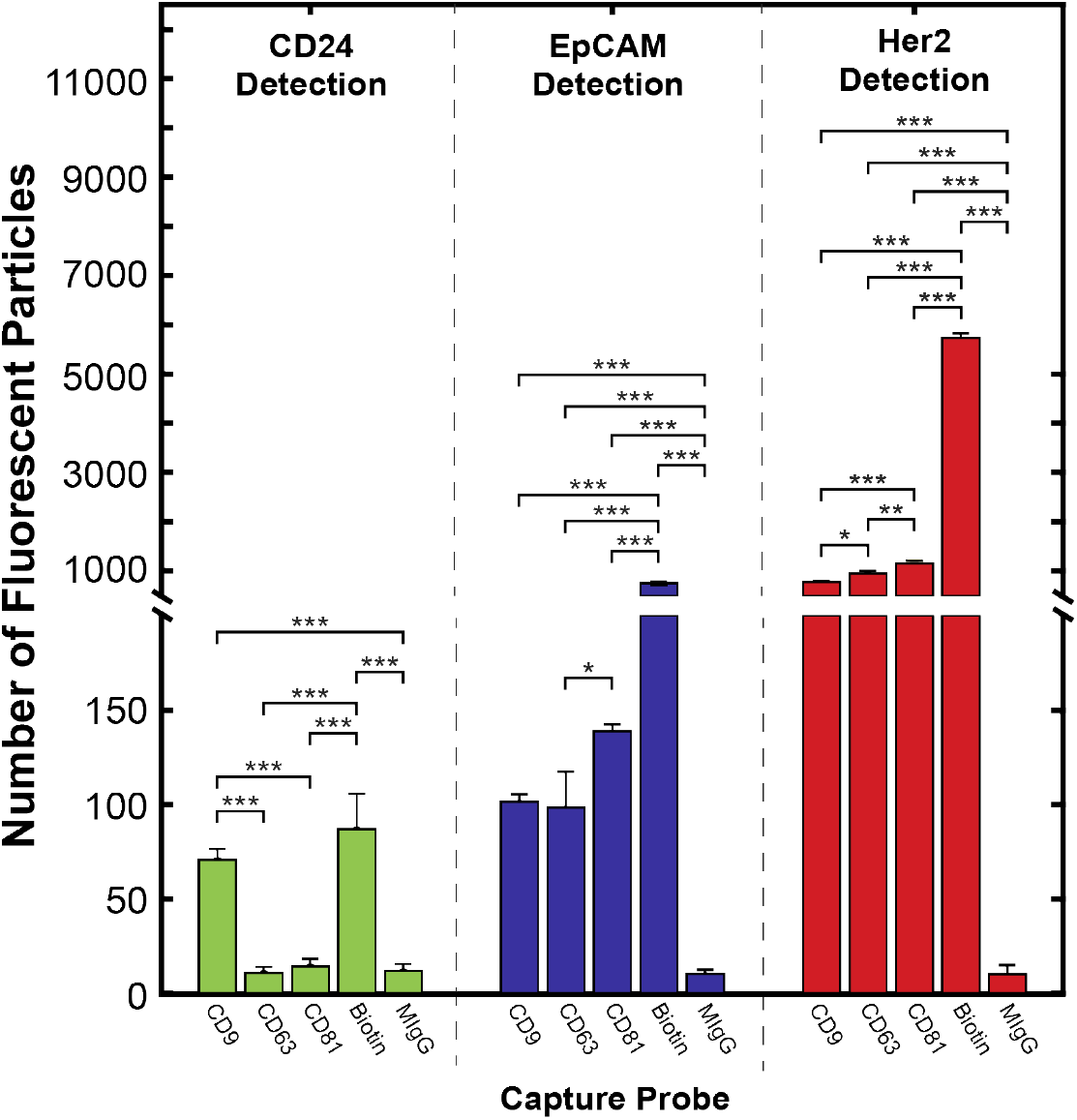
Biotinylated SK-OV-3 EVs were captured by anti-CD9, anti-CD63, anti-CD81, anti-biotin, or anti-MIgG, and detected by anti-CD24 (green), anti-EpCAM (blue), and anti-Her2 (red). CD24 is only multiplexed with CD9, while EpCAM and Her2 are associated with all three tetraspanins. Anti-biotin captured significantly more CD24+ and Her2+ EVs than any single tetraspanin. Error bars represent +/- one s.d. * p<0.05, ** p<0.01, *** p<0.001 by ANOVA.

CD24+ EVs were detected at a similar frequency on anti-CD9 and anti-biotin capture spots. However, anti-CD63 and anti-CD81 did not have a significant number of captured EVs above the MIgG control spot. Although EpCAM+ EVs could be detected on all three tetraspanin capture spots at similar rates, biotin capture identified more EVs by approximately a 10-fold difference. Her2+ EVs were identified with high frequency on all tetraspanin spots with approximately 5 times as many detected on the antibiotin capture spot.

## Discussion

Although EV-related research has grown exponentially in recent years, many basic questions remain, particularly related to compositional heterogeneity. Specifically, protein multiplex bias, which we define as the enrichment of EVs by a specific protein, may be limiting in the absence of accepted, universal exosome-exclusive markers. Understanding protein expression patterns across the true distribution of EVs will be vital to making informed decisions on use of EV capture for many applications.

### EVs from different sources have varying tetraspanin profiles

We isolated EVs from a variety of sources to determine single-EV tetraspanin colocalization patterns, including a cultured OvCa cell line (SK-OV-3), human serum, and cultured pMSCs. In all cases, we found that tetraspanin expression was not homogeneous across a single population of EVs, with varying number of EVs captured by each tetraspanin. In addition, we found that EVs from the same biofluid source, independent of isolation technique, had consistent tetraspanin colocalization profiles. However, this tetraspanin colocalization profile differed between EV sources, with SK-OV-3, pMSC, and serum derived EVs all exhibiting unique expression patterns. Serum EVs had the most similar profile to SK-OV-3 EVs but had a significantly higher percentage of anti-CD9 detected EVs on anti-CD9 and anti-CD63 capture spots. pMSC derived EVs had a much more unique tetraspanin profile with a higher anti-CD63 capture/anti-CD63 detection and anti-CD81 capture/anti-CD9 detection. In general, this supports that protein multiplex bias, even by a well-known EV associated marker, can impact the apparent protein expression within an EV population.

Here we corroborated previous reports that there are differences in tetraspanin profile between different EV sources^46,47,3^. Yet few have measured multiplexed tetraspanin expression. Kowal et al. used bead-based immunocapture and successive Western blot to examine multiplex expression of tetraspanins^12^. This work showed that CD9 and CD81 were found on most dendritic cell EVs but are not necessarily co-expressed. However, CD63 was on a much smaller subpopulation which was always co-expressed with either CD9 or CD81. Our data did support that CD63 was generally a less common marker, given that either CD9 or CD81 almost always captured more EVs. However, since the number of CD63+ EVs was not constant on each tetraspanin capture spot, we did not conclude that CD63 was always co-expressed with CD9 or CD81. This emphasizes the importance of understanding the specific profile of tetraspanins for a given EV source prior to utilizing any single tetraspanin for capture.

### Apparent tetraspanin profiles vary between SP-IRIS and flow cytometry

Currently, single-EV protein analysis techniques are hampered by limits of detection which severely restrict the population of EVs that can be analyzed. These techniques are generally either protein biased, requiring presence of a single protein for capture, or size biased, requiring a large enough size to be identified. While SP-IRIS, as performed here, is protein biased, requiring CD9, CD63, or CD81 to be present for analysis, the combination of these three EV-associated proteins helps to mitigate this limitation. On the other hand, flow cytometry, a useful technique for measuring single-EV protein expression, is scatter-biased. When the same EVs from each source were analyzed by flow cytometry, the acquired apparent tetraspanin profiles had variable agreement with those acquired by SP-IRIS, as described before. The lack of agreement between these two analysis techniques could be due to differences in sensitivity to the different fluorophores, as a result of un-bound antibody in the stream for flow cytometry, optics strength and collection efficiencies, or autofluorescence of the bound antibody. Although it is likely that this contributed to some of the differences in apparent tetraspanin profile, lack of major systematic bias, e.g., all “blue” fluorophores appearing more numerous, indicate it is only a minor effect. For example, anti-CD9 detection by flow cytometry decreased for serum EVs yet increased for pMSC EVs compared to immunocapture. Overall, using fluorescence-based detection, SP-IRIS is not limited by scatter of EVs, while flow cytometry can only see a small fraction of total EVs larger than 100 nm. This difference in the “visible” EVs that were analyzed may also be responsible for the difference in apparent tetraspanin expression. In one previous study, EV tetraspanin expression varied with apparent EV density, i.e., scattering intensity, subpopulations^12^. Specifically, CD63 was restricted to a less dense population of EVs. Since these results do not show a homogeneous decrease in CD63 detection by flow cytometry compared to immunocapture, it is difficult to tell if density is a major factor in our study. Another explanation could be that density-dependent tetraspanin expression is not homogeneous across cell sources. Size-dependent expression could also explain the changes; however, to our knowledge no group has systematically studied correlation between EV size and tetraspanin expression at single vesicle resolution. Nonetheless, the differences in apparent tetraspanin profile between immunocapture and flow cytometry emphasizes the importance of considering the technique specific limitations of EV analysis. In summary, flow cytometry may be more appropriate for high-throughput needs, while immunocapture/fluorescence likely provides a more wholistic protein expression due to its ability to visualize smaller particles in general.

### Non-specific EV capture decreases multiplexing bias in diagnostic application

There is increasing interest in using EVs as diagnostic tools due to their reflection of parent cell markers and their presence in multiple biofluids, decreasing invasiveness of diagnostic procedures. OvCa diagnosis is a prime candidate for next generation EV-screening since CA-125 ELISA, the current clinical gold standard, lacks both sensitivity and specificity, resulting in high rates of false-positives and overwhelmingly late-stage diagnosis^48^. Multiple groups have identified specific markers associated with OvCa on OvCa EVs including CD24, EpCAM, and Her2^43,44^. Currently, many cancer diagnostics rely on a primary capture antibody, often a tetraspanin, and use a secondary label for EV-associated markers^49,50,51^, including for OvCa^43^.

Although tetraspanins are well-known EV-associated markers, multiple groups have shown that these markers are not necessarily ubiquitous on EVs^46,47,3^, but that their expression changes with EV subclass, size, and source. We reasoned that capture by tetraspanin may bias downstream multiplexed marker detection, given our initial findings of high heterogeneity in tetraspanin expression for different sources of EVs. To explicitly test this, we chemically modified EVs to incorporate biotin at their surface, as a handle for non-specific capture by anti-biotin. Perhaps unsurprisingly, we found that anti-biotin was consistently able to capture at least as many or more EVs than any single tetraspanin. Unexpectedly, we found that SK-OV-3 EVs expressing CD24 were only multiplexed with CD9 (but not CD63 nor CD81). On the other hand, EpCAM and Her2 had a similar number of EVs captured by each tetraspanin. However, significantly more EpCAM and Her2 expressing SK-OV-3 EVs were detected when capturing non-specifically. This suggests that there may be a large percentage of non-tetraspanin expressing EVs that contain the bulk of these markers. Together these data show that single-tetraspanin capture of EVs can impact the efficiency and sensitivity of detection of these diagnostically relevant markers. This suggests that the multiplexing efficiency of disease markers with specific tetraspanins should be studied prior to choosing a single protein for capture. A combination of tetraspanins can be used to avoid bias or a non-specific capture method can be utilized as carried out here.

## Conclusion

The goal of this study was to determine the bias associated with single-tetraspanin capture of EVs. Here, we show that although EVs from a single source have consistent multiplexing patterns of tetraspanins regardless of isolation technique and passage, tetraspanins are not homogeneously expressed on every EV. This shows that EV subpopulations exist with unique tetraspanin density and multiplexing even from a single cell source, such that capture by a single tetraspanin type could inadvertently bias downstream profiling. Moreover, tetraspanin density and frequency was unique for different clinically relevant sources of EVs. This emphasizes that EV capture by any given antibody needs to be informed by careful characterization of the protein profile of a given EV source. Furthermore, we showed that non-specific EV capture by anti-biotin can be achieved by biotinylation of EVs. This allowed us to identify significant bias in apparent tetraspanin expression by single-tetraspanin captured EVs from both serum and pMSCs. Finally, when detecting OvCa markers on SK-OV-3 derived EVs, we identified that CD24 could only be identified by CD9 or non-specific capture, and that EpCAM and Her2 detection were more sensitive by non-specific capture. This work demonstrates that non-specific biotinylation may allow for less biased multiplexed analysis of EVs in diagnostics platforms and untangles some of the complex nature of EV protein expression. Careful consideration of EV-capture methods is necessary to maximize accurate and sensitive detection of multiplexed markers.

## Acknowledgements

R.P.C. was supported by a Research Scholar Grant, RSG-19-116-01-CDD, from the American Cancer Society. T.R. gratefully acknowledges The Sigrid Juselius Foundation, Helsinki, Finland. This work was supported by funds from the Ovarian Cancer Education and Research Network (OCERN), and the NIH (1R01CA241666). We also would like to thank members of the Wang Lab at University of California Davis Health for their work collected and de-identifying pMSCs for this work. Thanks to Joshua Welsh at the NIH for training in single particle flow cytometry techniques.

## Data availability

All raw data generated and analyzed during the current study are available in a Zenodo repository with the identifier doi:XX.XXX/XX (will be updated on acceptance).

