## Supplementary Information for "Tetraspanin immunocapture phenotypes extracellular vesicles according to biofluid source but may limit identification of multiplexed cancer biomarkers"

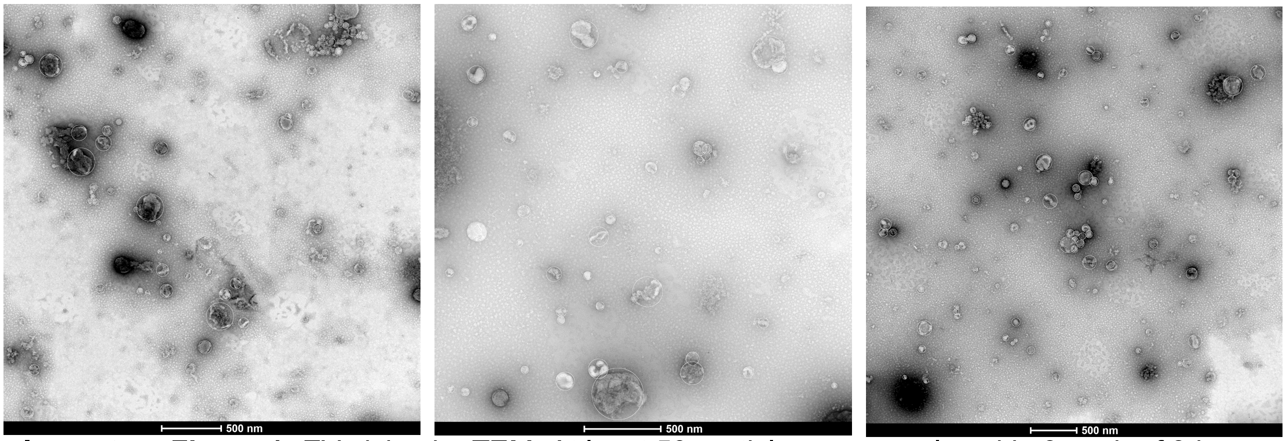


**Supplementary Figure 1:** EV sizing by TEM. At least 50 particles were analyzed in 3 each of 3 images taken of SK-OV-3 EVs isolated by UC. Diameter of individual particles was estimated in the Fiji package of ImageJ by circling deflated EVs, as seen by thin white lines overlayed on TEM images.

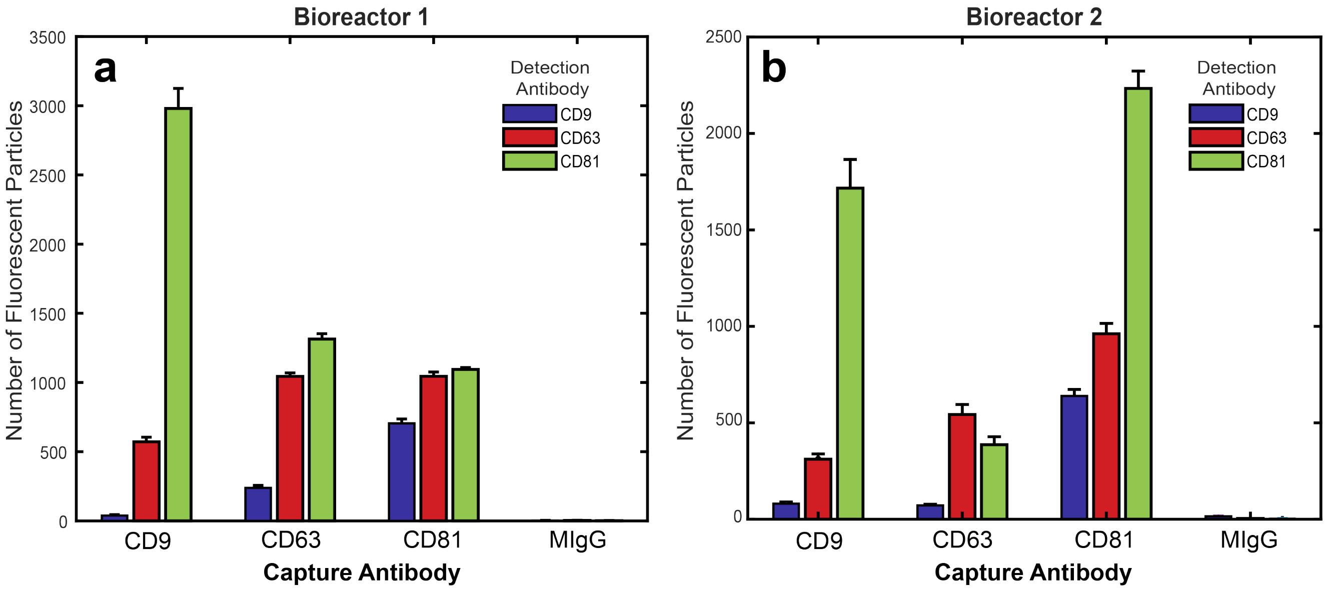


**Supplementary Figure 2:** SK-OV-3 EVs isolated from separate bioreactors. The tetraspanin profile of EVs from a first bioreactor (a), copied from Figure 3 for comparison, is similar to the tetraspanin profile of SK-OV-3 EVs isolated from a second bioreactor (b). Although there is a difference in CD81 capture/CD81 detection, this change was common between consecutive weeks from a single bioreactor.

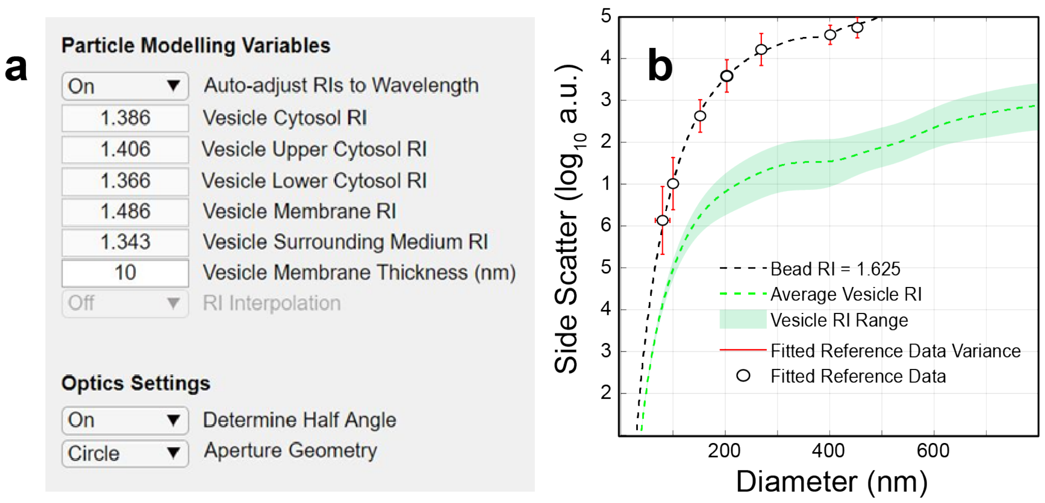


**Supplementary Figure 3:** $FCM_{PASS}$ Refractive Index Parameters and Modeling. Using scatter of a variety of standard beads, the size of SK-OV-3 EVs was modeled using (a) these parameters for average EV refractive index and membrane thickness. (b) $FCM_{PASS}$ model of SSC-H vs. diameter of EVs by measuring polystyrene bead SSC-H of varying sizes.

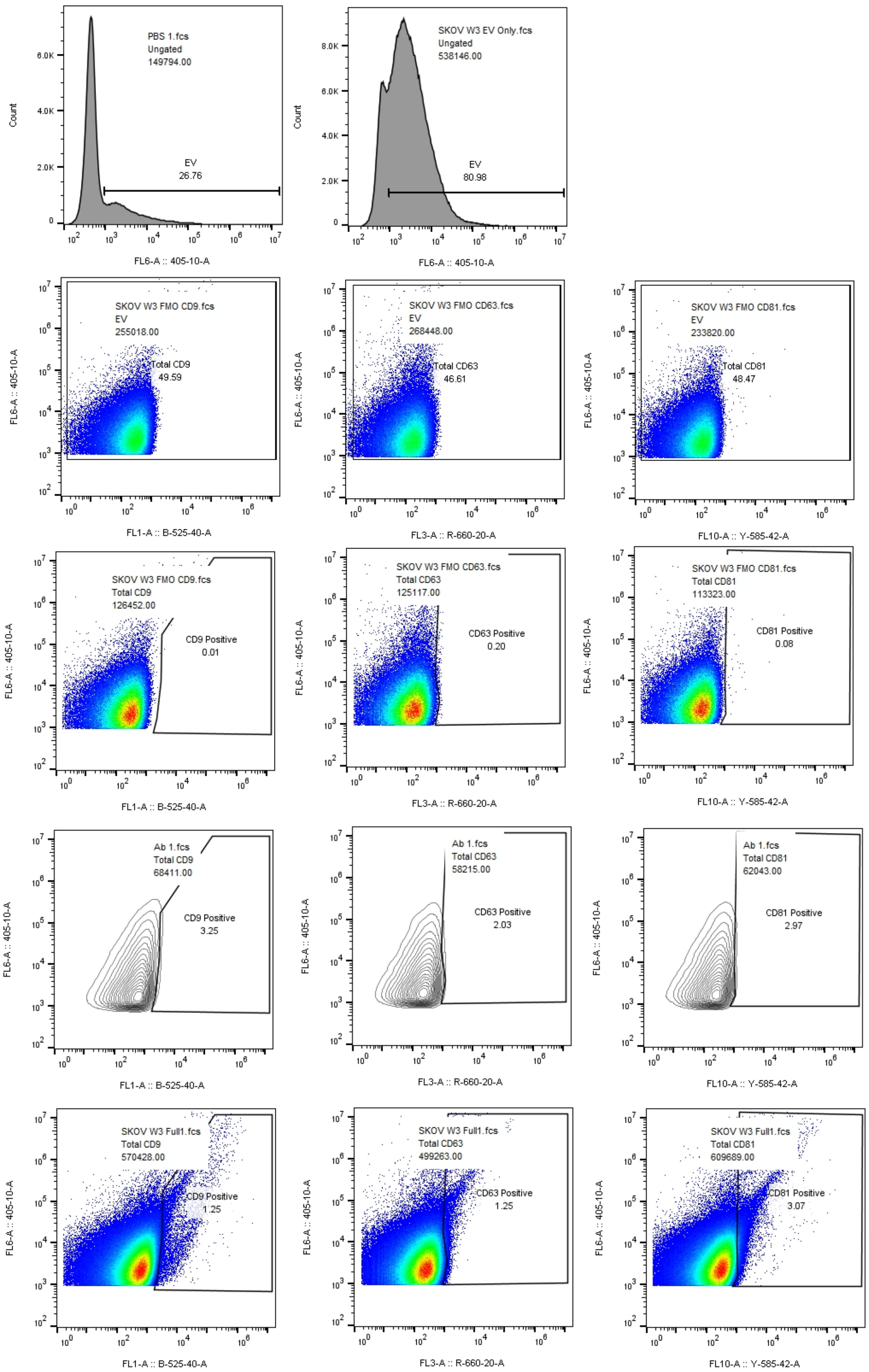


**Supplementary Figure 4:** Gating scheme for tetraspanin analysis. In row 1, EVs are gated from background noise by examining PBS versus an EV only control. In row 2, gates were used to separate the real events from those at the limits of fluorescence (at 0 fluorescence) to find the median fluorescent intensity of the EVs. In row 3, fluorescence-minus-one controls were used to gate fluorescent particles from unstained particles. In row 4 these gates were adjusted to encompass no more than ~5% of the antibody aggregates. In row 5, fully stained EVs are shown as an example of where these gates appeared on fluorescently labeled particles.

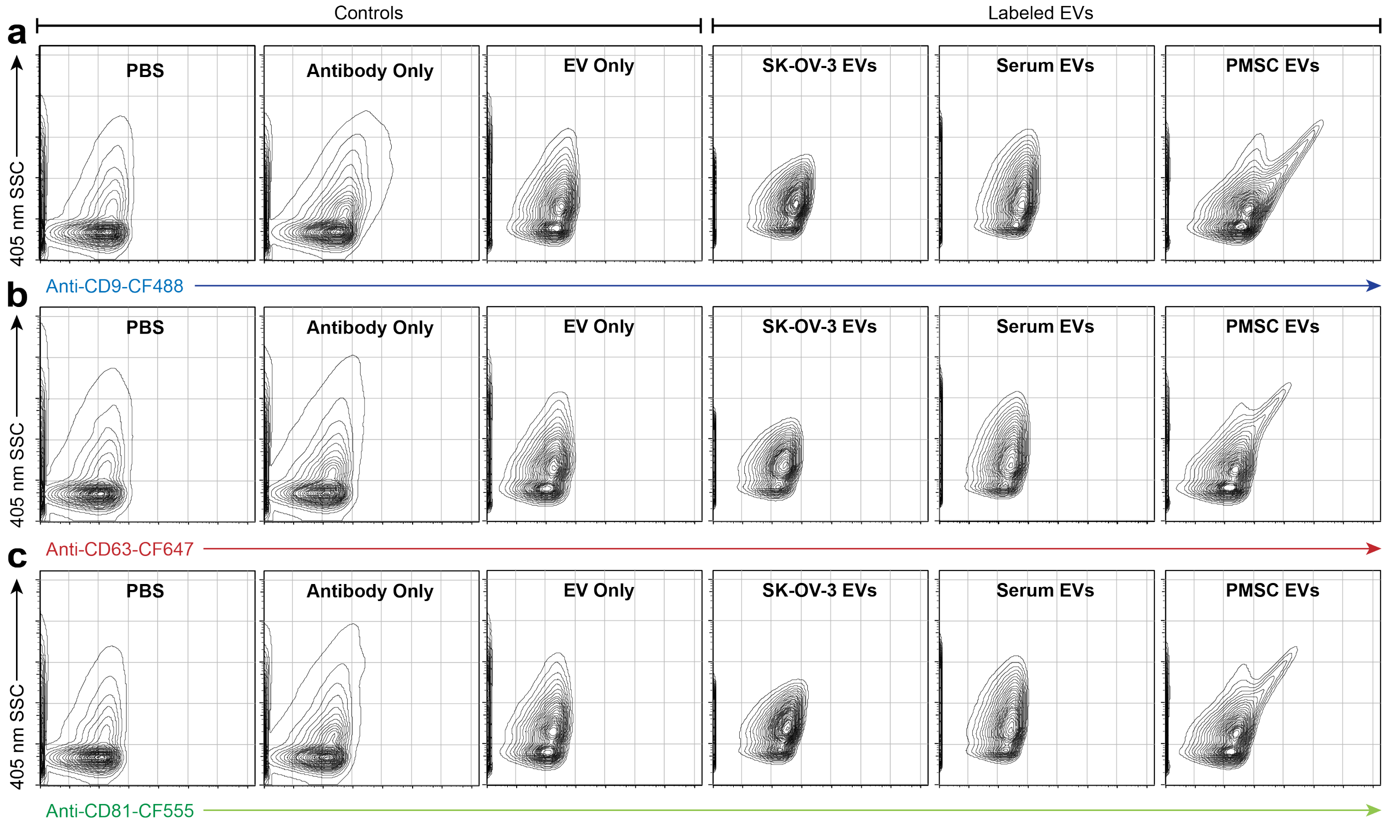


**Supplementary Figure 5:** Flow cytometry of EVs from various sources labeled with anti-tetraspanin antibodies. SSC-A vs (a) anti-CD9-CF488, (b) anti-CD63-CF647, and (c) anti-CD81-CF555 of control samples and labeled EVs. EV only sample is SK-OV-3 EVs.
